# Hippocampal engram networks for fear memory recruit new synapses and modify pre-existing synapses *in vivo*

**DOI:** 10.1101/2022.11.29.518458

**Authors:** Chaery Lee, Byung Hun Lee, Hyunsu Jung, Chiwoo Lee, Yongmin Sung, Hyopil Kim, Jooyoung Kim, Jae Youn Shim, Ji-il Kim, Dong Il Choi, Hye Yoon Park, Bong-Kiun Kaang

## Abstract

As basic units of neural networks, ensembles of synapses underlie cognitive functions like learning and memory. These synaptic engrams show elevated synaptic density among engram cells following contextual fear memory formation. Subsequent analysis of the CA3-CA1 engram synapse revealed larger spine sizes, as the synaptic connectivity correlated to the memory strength. Here, we elucidate the synapse dynamics between CA3 and CA1 by tracking identical synapses at multiple time points by adapting two-photon microscopy and dual-eGRASP technique *in vivo*. After memory formation, synaptic connections between engram populations are enhanced in conjunction with synaptogenesis within the hippocampal network. However, extinction learning specifically correlated with the disappearance of CA3 engram to CA1 engram (E-E) synapses. We observed “newly formed” synapses near pre-existing synapses, which clustered CA3-CA1 engram synapses after fear memory formation. Overall, we conclude that dynamics at CA3 to CA1 E-E synapses are key sites for modification during fear memory states.

**Highlights:**

- We adapted four-color two-photon microscopy with dual-eGRASP for a longitudinal observation of synaptic connections between CA3 and CA1.
- Synaptogenesis according to fear memory formation was specifically observed in E-E synapses.
- Extinction learning significantly correlated with the disappearance of E-E synapses.
- Particular distribution pattern of newly formed E-E synapses was observed after memory formation.

## Introduction

Memory engrams reflect the population of neurons activated during memory formation^1-3^. Reactivation or inactivation of these engram cells correlates to the recall or inhibition of the memory retrieval, respectively^1,3,4^. Evidence shows that the synapses of these engram cells physically encode the memory trace, implying that the synaptic networks between engram cells underlie memory formation, maintenance, and extinction^5-7^. Indeed, specifically enhanced synaptic connections between CA3 engram cells to CA1 engram cells were observed using “dual-eGRASP”^8^ that labels the pre- and post-synaptic part of GFP contacts^9,10^. The correlation of memory states and the synapses between engram cells reflect alterations in the spine morphology of activated neuronal ensembles^11^. Further, the localization of synapses within a dendritic branch is critical in memory formation throughout the brain^12-14^. Randomly scattered synapses primarily occur in sensory cortices, while synapses cluster in several other brain areas^12,13^, which may indicate hotspots of dendritic spines with enhanced turnover rates^14^. Whether such clustered spine formation is specific to engram synapses still remains an open question.

Technical limitations constrained prior studies to observations in different subjects at a specific memory state. Since the dual-eGRASP signals were obtained from perfused brain slices, longitudinal imaging within identical subjects was impossible. This technical approach hampered investigations on the dynamic changes in synaptic connectivity during various memory states *in vivo*. To overcome these issues, we integrated four-color *in vivo* two-photon imaging and dual-eGRASP system. Using this advanced approach, we sought to uncover the synaptic mechanisms that underlie engram synapse-specific enhanced connectivity during memory formation and the synaptic distributions on dendrites by comparing identical synapses across different memory states. Our analysis of synaptic dynamics between connected engram cells revealed: (1) higher proportion of “newly formed” synapses on postsynaptic engram dendrites, (2) a significant decrease in “E-E” synapses after memory extinction, and (3) a close distribution of “newly formed” synapses in sparsely innervated dendritic areas and clustering of “E-E” synapses.

## Results

### *In vivo* two-photon imaging enables longitudinal observation of dual-eGRASP in CA1

Using adeno-associated virus (AAV), we expressed dual-eGRASP^8^, which can label synapses with spectrally different fluorescence proteins (cyan and yellow) according to presynaptic motifs (Fig. 1A, left). To monitor random CA1 postsynaptic dendrites, we induced a sparse and constitutive expression of iRFP670 and post-eGRASP. To visualize CA1 engram dendrites at specific time points, we expressed myristoylated mScarlet-I driven by the Fos-dependent reverse tetracycline transactivator (Fos-rtTA)^15-18^. We induced a dense expression of cyan pre-eGRASP without Cre recombinase to cover most of the excitatory presynaptic inputs from contralateral CA3. To label synaptic inputs from a contralateral CA3 engram cell to CA1 neurons (Fig. 1A, middle), we expressed yellow pre-eGRASP using the Fos-rtTA system. These expression patterns enabled identifying CA3 engram to CA1 engram (E-E) synapses, which are recognized by yellow eGRASP signals on mScarlet-i expressing dendrites. Similarly, based on CA1 dendritic expression of mScarlet-i and presence of yellow GRASP on CA1 spines, we classified synapses into four classes between pre- and postsynaptic engram (E) and non-engram (N) cells: E-E, N-E, E-N, and N-N.

**Figure 1.**
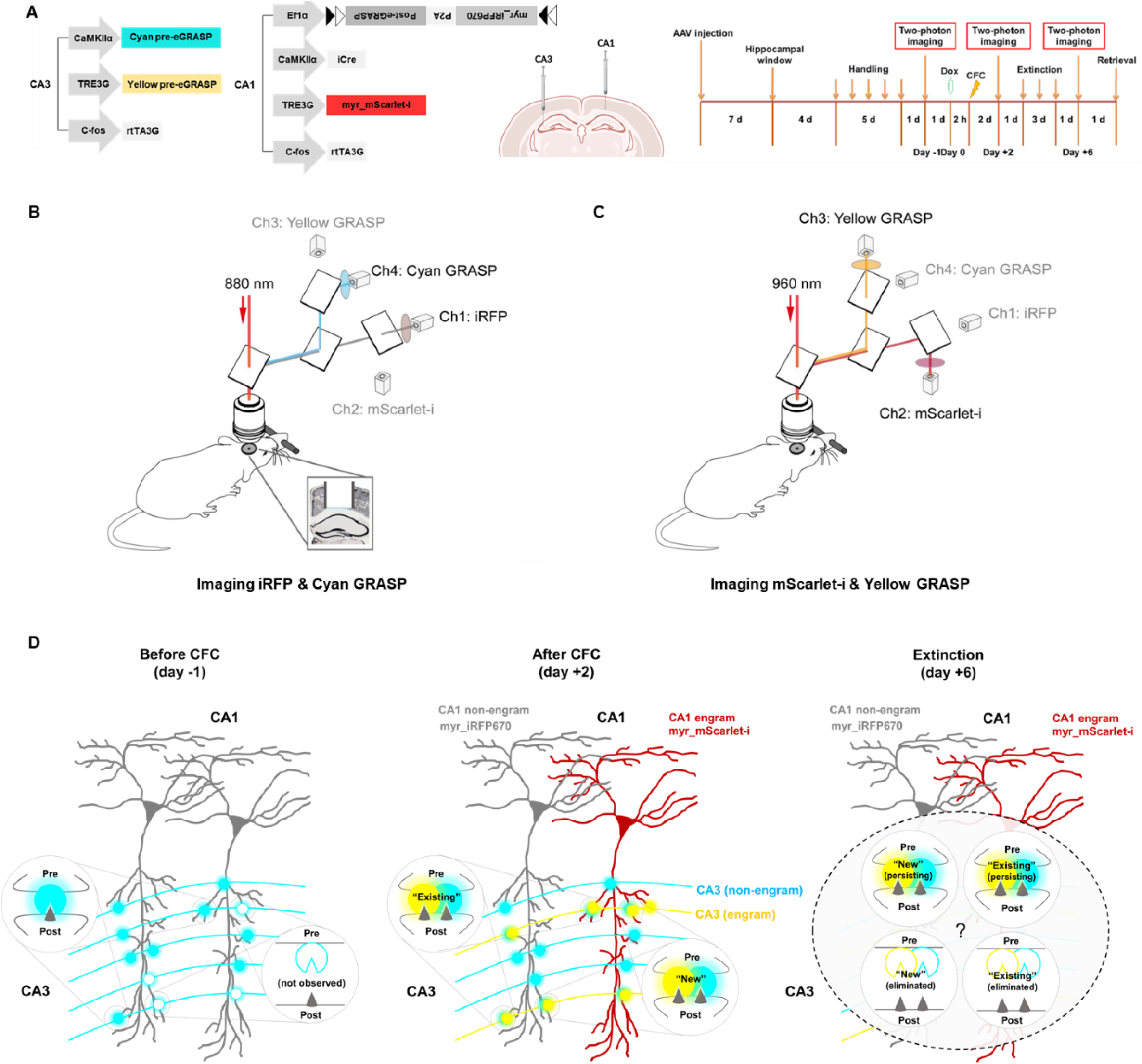
Experimental strategy diagrams to image synaptic connections *in vivo*. (A) (Left) Schematic illustration of AAVs injected into CA3 and CA1 of the hippocampus. (Middle) Virus cocktails were injected into contralateral CA3 and ipsilateral CA1 of the hippocampus. (Right) Schematic illustration of the experimental protocol to examine synaptic connections through memory formation and extinction. See also Figure S1. (B) Using an excitation wavelength of 880 nm, Cyan GRASP and random apical dendrites of CA1 (iRFP) were observed through the hippocampal window (inset). Fluorescence was collected via different PMTs after passing through dichroic mirrors and band-pass filters. (C) Yellow GRASP and apical dendrite of engram neurons in CA1 (mScarlet-i) were imaged using excitation wavelengths of 960 nm. (D) Schematic illustration of synapse classification based on their existence and persistence in each memory state.

After 7 days of recovery following AAV injection, hippocampal windows were implanted above dorsal CA1. After 14∼16 days of AAV expression, we conducted *in vivo* two-photon imaging of dual-eGRASP signals in the mouse CA1 before and after contextual fear conditioning (CFC) and after the fear extinction (Fig. 1A, right and Fig. S2A). Four types of fluorescence proteins (cyan eGRASP, yellow eGRASP, mScarlet-i, and iRFP670) were imaged with an excitation laser tuned at two different wavelengths (Fig. 1B, C).

Identical CA1 pyramidal cell dendrites were tracked by two-photon imaging in multiple areas of CA1 (Fig. 1D), and the images were further processed to reduce motion artifacts and to improve the resolution by deconvolution. Before fear conditioning, constitutively expressed cyan eGRASP covered most of the existing excitatory synapses in the CA3 to CA1 circuit. Thus, in terms of synapse dynamics, we could investigate whether a synapse was newly formed after fear conditioning or already existed by comparing the cyan eGRASP signals before CFC, after CFC, and after fear extinction. Combining this turnover information with engram synapse classification, we could map the persistence of engram synapses upon fear conditioning and extinction. Further, we could reveal if the increased synaptic density of E-E synapses derived from “newly formed” or “existing” synapses. In addition, we could analyze if each synapse was persistent or eliminated using the two-photon images taken after fear extinction.

### New synapses form primarily at E-E connections during memory formation

From the two-photon images of mouse CA1, we observed four kinds of synapses formed between CA3 and CA1 non-engram or engram (Fig. 2A, B). “Cyan only” synapses on the non-engram dendrites labeled only by iRFP670 represented N-N, while those on the engram dendrites that were labeled with both iRFP670 and mScarlet-I represented N-E. Moreover, “yellow only” and “cyan + yellow” synapses on the non-engram dendrites represented E-N, while those on the engram dendrites represented E-E. Only few synapses were labeled as “yellow only” after memory formation, suggesting that cyan eGRASP already covered most of the existing synapses. By manually comparing the dual-eGRASP signals of these four types of synapses at our three time points (before or after memory formation, and after fear extinction), we could investigate the synaptic dynamics in more detail.

**Figure 2.**
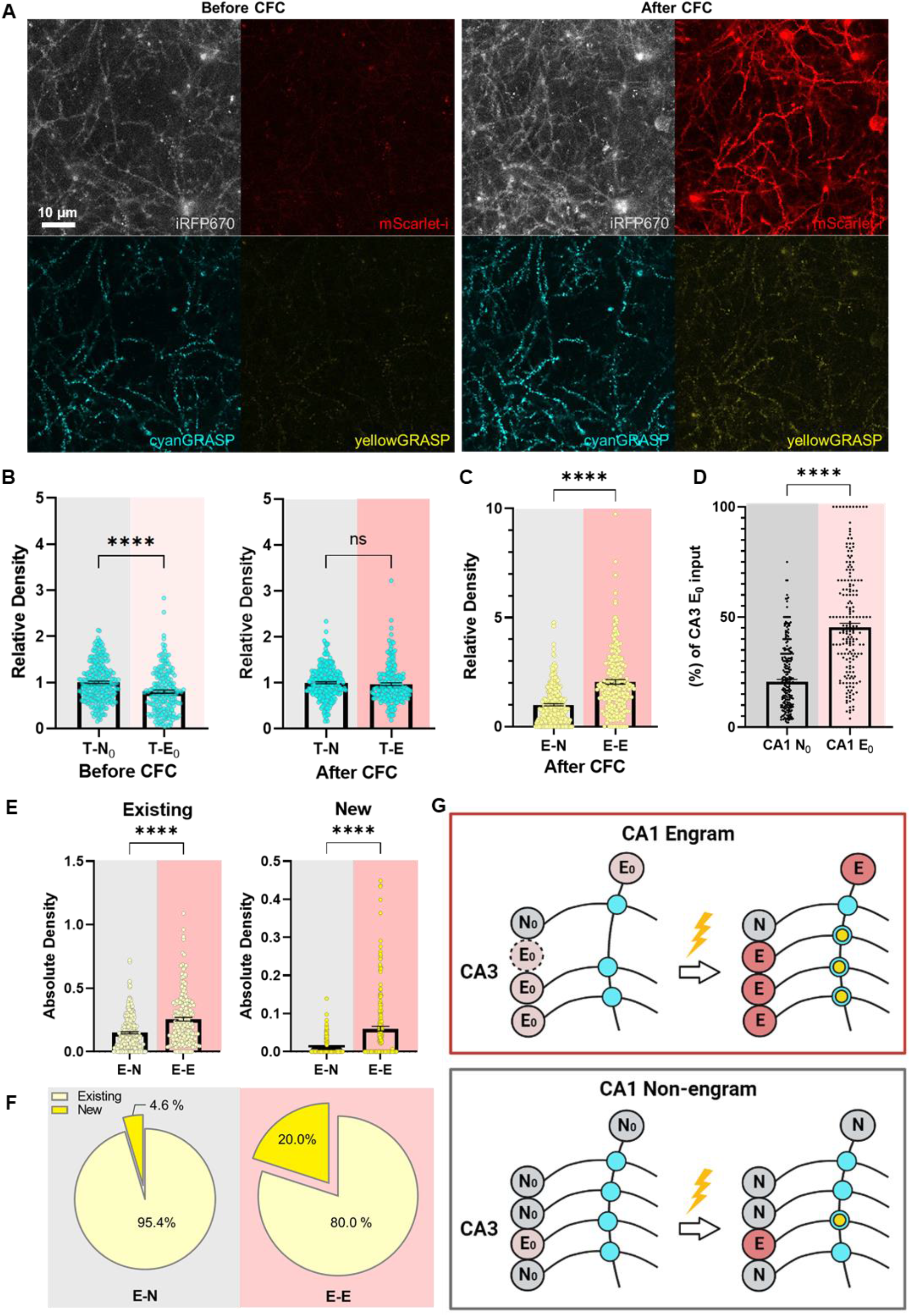
Elevated “existing” and “newly formed” synaptic populations in CA1 engram dendrites after CFC. (A) Representative two-photon images of dual-eGRASP signals in a non-engram or engram dendrite before and after CFC. (B) Relative density of random cyan GRASP on non-engram or engram dendrites before and after fear conditioning; CA1 non-engram dendrites (n = 199); CA1 engram dendrites (n = 266); Mann-Whitney two-tailed test. (Left, Before CFC) ****P < 0.0001. (Right, After CFC) P = 0.4377. N_0_ and E_0_ indicate CA1 cells before CFC that become non-engram and engram later after CFC, respectively. (C) Relative density of yellow GRASP on non-engram or engram dendrites after fear conditioning; CA1 non-engram dendrites (n = 199); CA1 engram dendrites (n = 266); Mann-Whitney two-tailed test. **** P < 0.0001. (D) Dendritic proportion of synaptic inputs from CA3 E_0_ before CFC. The ratio of particular cyan eGRASP, tracked as cyan+yellow eGRASP after CFC among total cyan eGRASP, was measured; CA1 N_0_ dendrites (n = 199); CA1 E_0_ dendrites (n = 266); Mann-Whitney two-tailed test. **** P < 0.0001. (E) The absolute densities of “existing” or “newly formed” synapses. CA1 non-engram dendrites (n = 199); CA1 engram dendrites (n = 266); (Left) Mann-Whitney two-tailed test. ***P < 0.0001; (Right) Mann-Whitney two-tailed test. ***P < 0.0001. (F) The proportion of “existing” and “newly formed” synapses in non-engram or engram dendrites; “existing” N-E synapses (n = 1122); “new” N-E synapses (n = 54); “existing” E-E synapses (n = 1245); “new” E-E synapses (n = 311); CA1 non-engram dendrites (n = 199); CA1 engram dendrites (n = 266). (G) Schematic illustration of the overall synaptic dynamics between CA3 and CA1 neurons before and after CFC. (Upper box) CA1 engram; (Lower box) CA1 non-engram. Circles entitled as N_0_ and E_0_ indicate CA3 and CA1 neurons that were each labelled as non-engram and engram after CFC, respectively. E_0_ with a dotted circle indicates CA3 neuron whose presence remains unknown. Circles on crossed lines indicate cyanGRASP or cyan & yellowGRASP. Data are represented as mean ± SEM in all figures.

We first observed an elevated relative synaptic density of E-E connections compared to E-N after fear memory formation, reproducing our previous report^8^ (Fig. 2C). The synaptic density of total presynaptic cells to engram (T-E) was significantly lower than T-N before fear conditioning (Fig. 2B, left). However, no differences between T-N and T-E were found after memory formation, consistent with the previous data^8^ (Fig. 2B, right and Fig. S1). These confirmatory data indicate that engram synapses preferentially form on dendrites with lower density of synaptic inputs from CA3. Our advanced methodology for *in vivo* eGRASP did not significantly impact our previous results. We next examined the dendritic proportion of synaptic inputs that CA1 dendrites receive from CA3 cells (E_0_) before fear conditioning that eventually become engrams after fear conditioning. CA1 E_0_ dendrites received a significantly higher percentage of inputs from CA3 E_0_ neurons compared to N_0_ dendrites that still remained non-engrams after fear conditioning (Fig. 2D). Moreover, to elucidate whether the increased synaptic density derived from “existing” or “newly formed” synapses, we tracked the history of dual-eGRASP signals of synapses on either non-engram or engram dendrites labeled as “cyan + yellow” after memory formation. Interestingly, both “existing” and “newly formed” synapses accounted for the elevated synaptic density of the E-E connection (Fig. 2E). However, the proportion of “newly formed” synapses was significantly higher in engram dendrites compared to non-engram dendrites, indicating that “newly formed” synapses may exert a greater contribution for memory formation at the synaptic level in memory engram networks (Fig. 2F). Such synaptic dynamics were not observed when the mice only underwent context exposure without any conditioning (Fig. S3). In conclusion, CA1 engram dendrites are preferentially connected with CA3 engram cells even before fear conditioning, while CA1 non-engram dendrites receive unbiased inputs from CA3 neurons. Interestingly, some neurons with dendrites receiving less excitatory synapses were recruited as engram dendrites due to the major input from CA3 potential engram cells (Fig. 2G).

### “E-E” synapses significantly decrease after memory extinction

To examine changes at the synaptic level according to memory attenuation, we tracked identical spines after fear memory extinction (Fig. S2C, D). Particularly, we observed the dynamics of synapses labeled as “cyan + yellow” after contextual fear conditioning on either CA1 engram or non-engram cells. Dendrites that were previously selected for analysis after memory formation were analyzed again, and we determined whether each dual-eGRASP signal remained or completely disappeared from the two-photon images (Fig. 3A). Since cyan-GRASP was constitutively expressed during the experiment, some of the “cyan only” synapses were also generated or disappeared after fear extinction. So, we tracked “total cyan” GRASP signals, which included both “cyan only” and “cyan + yellow”, during the entire experiment. The relative spine density of total cyan GRASP synapses was significantly increased after fear memory formation, while we could not observe any significant changes after fear extinction (Fig.S1).

**Figure 3.**
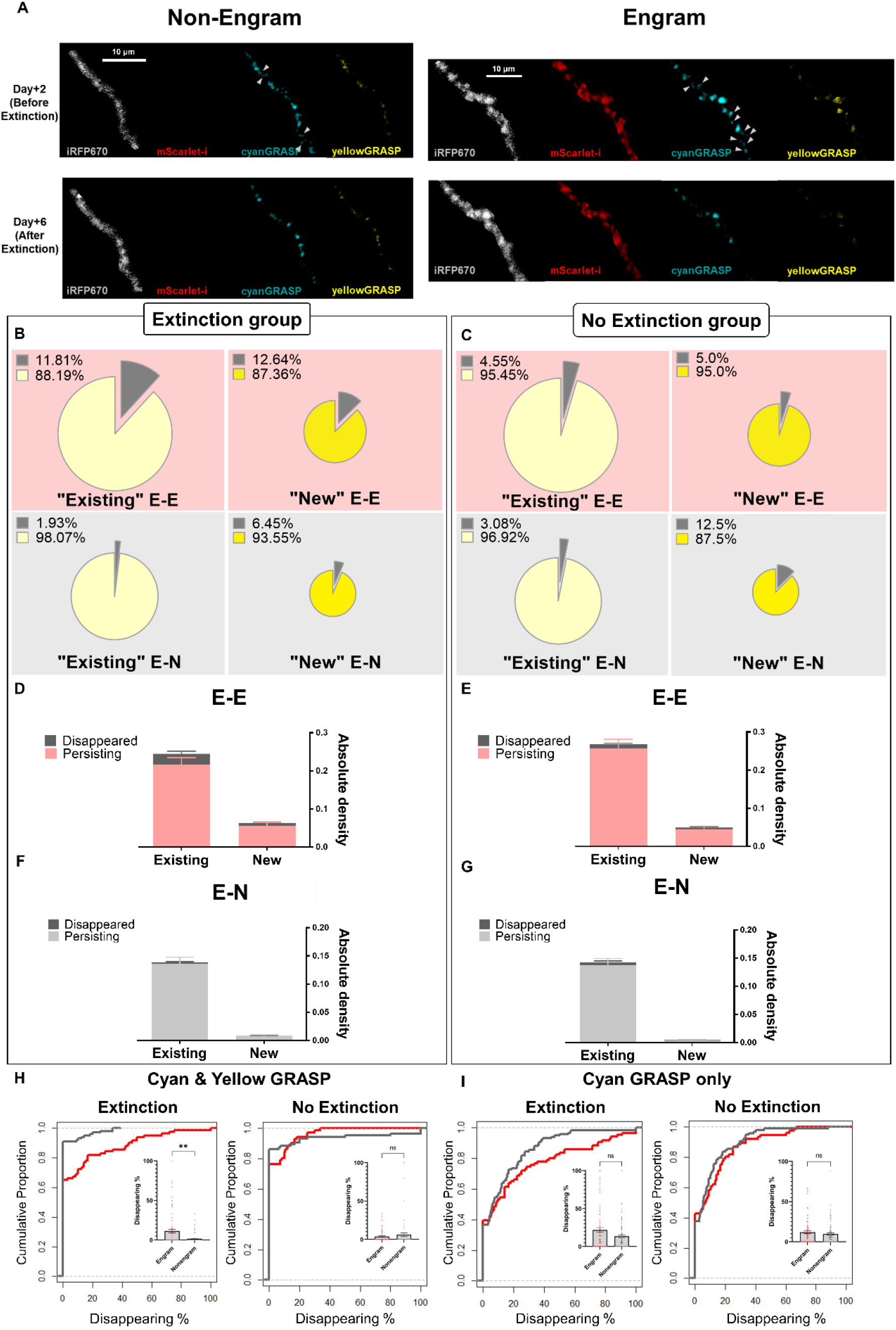
Extinction eliminates both existing and new engram-engram synapses. (A) Representative image of engram and non-engram dendrites after fear extinction training. Arrows indicate dual-eGRASP signals that disappeared after fear extinction. See also Figure S3. (B) Proportions of persisting and disappearing engram synapses in the extinction group. Yellow and gray indicate persisting and disappeared percentages of the synapses, respectively. Disappeared “existing” E-E synapses (n = 75); persisting “existing” E-E synapses (n = 560); disappeared “new” E-E synapses (n = 22); persisting “new” E-E synapses (n = 152); disappeared “existing” E-N synapses (n = 10); persisting “existing” E-N synapses (n = 507); disappeared “new” E-N synapses (n = 2); persisting “new” E-N synapses (n = 29). (C) Proportions of persisting and disappearing engram synapses in the no extinction group. Yellow and gray indicate persisting and disappeared percentages of the synapses, respectively. Disappeared “existing” E-E synapses (n = 23); persisting “existing” E-E synapses (n = 482); disappeared “new” E-E synapses (n = 5); persisting “new” E-E synapses (n = 95); disappeared “existing” E-N synapses (n = 13); persisting “existing” E-N synapses (n = 409); disappeared “new” E-N synapses (n = 2); persisting “new” E-N synapses (n = 14). (D) Absolute synaptic density of existing and new E-E synapses in the extinction group. CA1 engram dendrites (n = 86); CA1 non-engram dendrites (n = 117). (E) Composition of existing and new E-E synapses in the no extinction group. CA1 engram dendrites (n = 78); CA1 non-engram dendrites (n = 98). (F) Absolute synaptic density of existing and new E-N synapses in the extinction group. CA1 engram dendrites (n = 86); CA1 non-engram dendrites (n = 117). (G) Absolute synaptic density of existing and new E-N synapses in the no extinction group. CA1 engram dendrites (n = 78); CA1 non-engram dendrites (n = 98). (H) Cumulative plot of disappearing percentage of E-E or E-N synapses on CA1 dendrites of extinction and no extinction group. CA1 engram dendrites of extinction group (n = 86); CA1 non-engram dendrites of extinction group (n = 117); CA1 engram dendrites of no extinction group (n = 78); CA1 non-engram dendrites of no extinction group (n = 98); Kolmogorov-Smirnov test, Extinction, p = 0.0039; No Extinction, p = 0.7305. (I) Cumulative plot of disappearing percentage of N-E or N-N synapses on CA1 dendrites of extinction and no extinction group. CA1 engram dendrites of extinction group (n = 86); CA1 non-engram dendrites of extinction group (n = 117); CA1 engram dendrites of no extinction group (n = 78); CA1 non-engram dendrites of no extinction group (n = 98); Kolmogorov-Smirnov test, Extinction, p = 0.1989; No Extinction, p = 0.4428. Data are represented as mean ± SEM in all figures.

From our analysis of two-photon images taken after fear extinction, we first observed the disappearance of either “New” or “Existing” E-E synapses, which both occurred approximately at a 12% frequency (Fig. 3B). Compared to E-N synapses, a significantly higher percentage of “existing” E-E synapses disappeared (Fig. S4). Interestingly, the decrease of absolute spine density due to the disappearance of some synapses was limited to E-E synapses, while nearly none of the E-N synapses disappeared (Fig. 3D, F). Such E-E specific disappearance of “existing” synapses occurred only when the subjects underwent fear extinction, as E-E and E-N synapses showed insignificant difference in the no extinction group (Fig. 3C, E, G). Disappearance of synapses from CA3 engrams was biased to CA1 engram dendrites by extinction training, while those from CA3 non-engrams did not show any significant differences (Fig. 3H, I). Repeated context exposure did not induce any particular disappearance of synapses (Fig. S3F, G, H). Based on these data, we propose that contextual fear memory is highly correlated with the alteration of CA3-CA1 E-E synapses during memory attenuation.

### Spatial distribution of “newly formed E-E” synapses cluster E-E synapses

Synapses are distributed in a scattered pattern on dendrites, which is determined by their anatomical structures^19^. However, another proposal claims that the clustering of dendritic spines with various spine morphologies also pre-determines the general location of synapses^20^. Here, we examined whether the newly formed spines elicited by CFC showed particular pattern of spatial distribution. First, we analyzed the dendritic spines labeled with dual-eGRASP at a spine level (Fig. 4A). Given the longitudinal images of synapses and dendrites, we could identify the 3-dimensional coordinates of synapses and measure the distance between each class of synapses. Even though both E-E and N-E synapses were newly formed after memory formation, only the mean distance between “new” E-E synapses was significantly shorter than the overall average distance of random synapses (Fig. 4B). This indicated that the “new” E-E synapses are closely formed with each other compared to other types of synapses.

**Figure 4.**
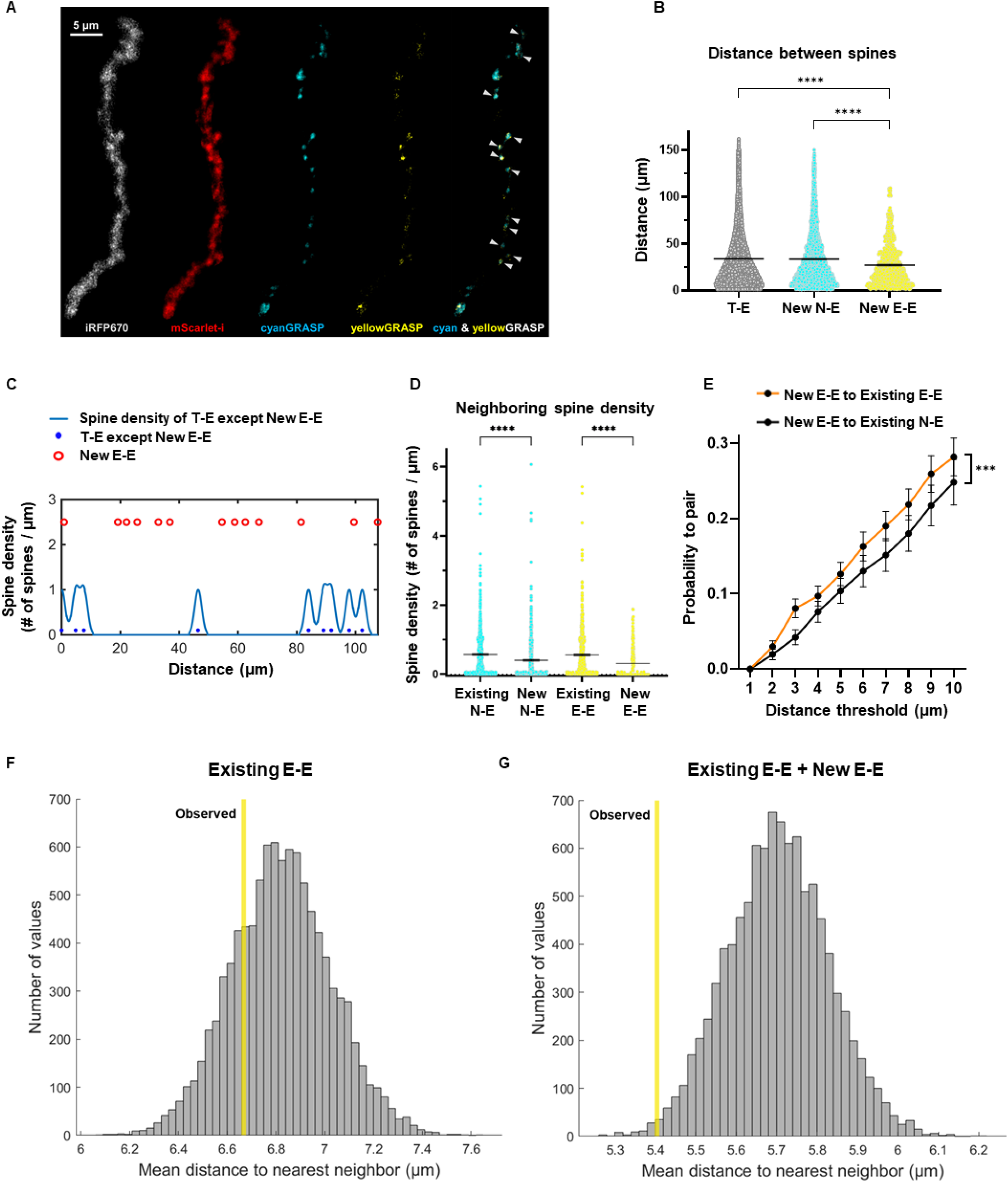
Dendritic distribution of synaptic connections among engram cells. (A) Representative image of the spatial distribution of newly formed E-E synapses on CA1 engram dendrite. Arrows indicate newly formed E-E synapses. (B) Distances measured between spines within identical categories. Each dot indicates distance measured between two random spines within a dendrite segment. Pairs of “new” E-E synapses (n = 952); pairs of “new” N-E synapses (n = 5262); pairs of T-E synapses (n = 66436); Dunn’s multiple comparisons test after Kruskal-Wallis test. Kruskal-Wallis: ****P < 0.0001; Dunn’s ****P < 0.0001. (C) Spine density histogram (light blue) and one-dimensional plotted spines (blue) / newly formed E-E spines (red) on a representative dendrite. (D) Spine densities (Spine number / 2.5 μm) measured at the dendritic locations of four different types of synapses. Each dot indicates surrounding spine density that was measured by the number of spines within a 2.5 μm-wide moving window. Density of surrounding “pre-existing” N-E and E-E synapses was calculated. “Existing” N-E synapses (n = 1,003); “new” N-E synapses (n = 655); “existing” E-E synapses (n = 841); “new” E-E synapses (n = 230); Dunn’s multiple comparisons test after Kruskal-Wallis test. Kruskal-Wallis: ****P < 0.0001; Dunn’s ****P < 0.0001. Data are represented as mean ± SEM in all figures. (E) The number of pairs between two groups of synapses existing within various distance thresholds divided by the total number of possible pairs selected between two groups of synapses. “New E-E to Existing E-E” indicates the pairs between ‘newly formed’ E-E synapses and ‘existing’ E-E synapses (n = 76); “New E-E to Existing N-E” indicates the pairs between ‘newly formed’ E-E synapses and ‘existing’ N-E synapses (n = 69); Two-way ANOVA; Distance threshold, F(9, 1430) = 47.29, ****P < 0.0001; Type of synapse pair, F(1,1430) = 11.43, ***P < 0.001; Interaction, F(9, 1430) = 0.2900, P = 0.9777. (F, G) Null distribution of the average nearest neighbor distance (NND) from “existing” E-E synapses (F) and total E-E synapses (G). 10,000 simulations of randomized label of spines were run, while the coordinates and number of each label of spines were controlled. Yellow line indicates the actual average NND (6.67 μm in existing E-E synapses, 5.40 μm in total E-E synapses) observed in the data.

We further visualized the distribution of “existing” and “newly formed” synapses by projecting the dual-eGRASP signals on a one-dimensional graph. We found that ‘new’ synapses (red circles in Fig. 4C) were closely formed in some dendritic regions with significantly lower spine density (Fig. 4C, D). The E-E synapses generated after fear memory formation tended to locate on sparser dendritic areas with higher capacity. “Newly formed” E-E synapses were significantly paired with “existing” E-E synapses rather than with “existing” N-E synapses (Fig. 4E). Thus, E-E synapses form near existing E-E synapses after fear memory formation. Moreover, the average nearest-neighbor distance (NND) of total E-E synapses was significantly smaller than random chance (Fig. 4G), while that of “existing” E-E synapses was comparable to chance level (Fig. 4F). Such data indicate that the formation of “new” E-E synapses near dispersed “existing” E-E synapses resulted in the clustering of E-E synapses after memory formation. Such learning-related distribution of spines is known to occur within dendritic segments with increased spine turnover, which also increases network sparsity and memory capacity in the retrosplenial cortex^13^. Thus, this data indicates that new E-E synapses could also form within the dendritic hotspots.

## Discussion

Here, we adapted two-photon imaging with the dual-eGRASP system, enabling us to track the same synapse on specific dendrites over time *in vivo*. Since two-photon imaging permitted measurements at multiple time points, we could analyze the changing synaptic dynamics occurring in different memory states induced by fear memory formation or extinction. Engram-specific features at a synaptic level have been previously researched by adapting the dual-eGRASP system. Spine density and morphologies at synaptic connections between CA3 engrams and CA1 engrams significantly increase after fear memory formation^8^. However, limitations with confocal imaging using the dual-eGRASP system necessitated using a single animal and restricted time points to visualize the synapses. Since the data were obtained from different individuals, observing the actual changes occurring in an identical subject *in vivo* was impossible. Our newly developed method overcomes these challenges.

Since motor or auditory fear memory formation can generate new spines^21-23^, we predicted that “newly formed” synapses would underlie the elevated synaptic density at the E-E synaptic connection. We found that “newly formed” synapses accounted for a relatively high proportion of the elevated synaptic density in CA1 engram dendrites. Yet, we also found that the absolute spine density of “existing” synapses was also significantly higher in CA1 engram dendrites. Thus, memory formation requires both new and modified pre-existing synapses. We propose that “newly formed” synapses support the modifications to “existing” synapses to expand engram memory networks. This then raises the question of which cells have the potential to become engram cells. Based on our data, we posit that cells with lower spine density and higher memory capacity have a higher probability to become engram cells by forming new synapses and possibly increasing the neuronal inputs from the connections and excitability. Surprisingly, we found that such cells from CA3 and CA1 appeared to be connected to each other even before memory formation unlike the allocation theory in which neurons with fluctuating excitability are recruited as engram. This data implicated that recruitment for memory trace might be rather based on the pre-existing connectivity between neurons.

Two mechanisms have been proposed to underlie fear memory extinction - ‘unlearning’ of pre-acquired memories and ‘new learning’ about the contingency^24,25^. A recent study demonstrated that newly formed spines in the auditory cortex during the auditory fear conditioning were eliminated upon extinction of that memory^26^. Specifically, the enhanced spine morphology in the lateral amygdala was weakened following fear memory extinction^11^. While our previous study focused on the changes of spine morphology in the auditory cortex to lateral amygdala circuit, we examined the dynamics of synaptic density in the hippocampus by using our fear conditioning paradigm and new method. We found that some E-E synapses were eliminated following fear memory extinction. These data would support the ‘unlearning’ hypothesis. Since both “newly formed” and “existing” synapses are vulnerable to memory extinction, we suspect that these E-E synapses within the dorsal hippocampus are correlated with fear memory traces. By utilizing our experimental scheme on various brain regions, whether the mechanism is universal could be further elucidated.

Finally, our data indicate that “newly formed” synapses are closely distributed in particular dendritic areas with lower spine density. Although the distance between these “new” E-E synapses were relatively shorter, the distance between other “existing” E-E synapses or “cyan only” N-E synapses were relatively longer. In addition, “newly formed” E-E synapses were closely distributed with “existing” E-E synapses. New spines in motor cortex form during learning nearby pre-existing spines that exhibited task-related activity ^27^. Since longitudinal two-photon imaging enabled distinguishing “new” E-E synapses from “existing” E-E synapses, we speculated that such learning-related events also occur at the E-E synapse level. Thus, we observed how learning induces the clustering of synaptic engrams in the dorsal hippocampus *in vivo*.

Clustering of spines is well known to affect physiological properties in cortical areas. For instance, by enabling synaptic tagging and capturing, clustered plasticity is known to convert E-LTP/E-LTD to L-LTP/L-LTD^28^. Moreover, synaptic plasticity mediated by NMDA receptor is crucial for hippocampal spatial memory^29^. Clustered synapses are presumed to induce local dendritic spikes mediated by NMDA receptors, even in the absence of somatic action potential firing^30^. Such correlation between spine clustering and biological properties has also been observed in the hippocampus. For instance, structural plasticity of L-LTP was inversely proportional to the distance between dendritic spines within CA1 *ex vivo* slices^31^. Plasticity related protein products (PrPs) were highly shared between spines with shorter distance less than 50 μm^31^. Thus, shorter mean distance between “newly formed” synaptic engrams in Fig. 4B implies that those synapses contribute to clustered plasticity. We posit that new hippocampal memory clusters CA3 to CA1 E-E synapses after incorporating synapses that existed before learning. However, further studies should elucidate the functional and physiological relevance of clustering of “newly formed” synapses *in vivo*.

In this study, we also encountered several limitations in this *in vivo* dual-eGRASP approach. First, color schemes for the labeling of CA1 engram and non-engram dendrites were limited since two channels of two-photon microscopy were occupied by cyanGRASP and yellowGRASP. Such technical limitation led us to use the constitutive expression of iRFP670 in random CA1 dendrites. On our current two-photon microscopy setup, iRFP670 appeared to be extremely blur and vulnerable to bleaching. Thus, it was difficult to analyze additional data related to spine morphology such as spine size or volume from CA1 dendrites. Moreover, tracking dendritic spines in real-time via *in vivo* live imaging was hindered due to the motions caused by breathing of mice during imaging. Motion correction and image processing were required for GRASP analysis even though two-photon images were acquired from anesthetized mice. Further studies with two-photon imaging of awake mice would require advanced image stabilization for spine-level analysis. Moreover, additional studies will be required for functional manipulation of E-E synapses. Revealing the causal relation between the dynamics of specific spines and learning and memory still remains as a challenge.

Despite the limits, we elucidated the synaptic dynamics in CA1 by combining longitudinal *in vivo* two photon imaging with dual-eGRASP for the first time. Our findings successfully advance previous studies by enabling the classification of E-E synapses according to their existence before fear memory formation. Thus, we demonstrate that synaptic connections between engram cells specifically accompany synaptogenesis. These connections are also preferentially influenced by memory extinction. Furthermore, theoretical studies have proposed that clustered plasticity may be crucial for memory storage. Our data imply that clustering may occur at interregional engram populations that undergo synaptogenesis induced by memory formation. In conclusion, our results strongly support the hypothesis that Schaffer collateral E-E synapses are the physical substrates of fear memory traces.

## Supporting information

Key resource table

## STAR★Methods

### Resource availability

#### Lead contact

Further information and requests for resources and reagents should be directed to and will be fulfilled by the lead contact, Bong-Kiun Kaang.

#### Data and code availability

All data to understand and assess the conclusions of this study are available in the main text or supplementary materials. The original datasets generated during or analyzed during the current study are available from the corresponding authors.

#### Materials availability

All the materials used in this study are available upon request.

## Acknowledgements

This work was supported by the National Honor Scientist Program (NRF-2012R1A3A1050385) of Korea. B.H.L., J.Y.S., and H.Y.P. were supported by the Samsung Science and Technology Foundation under Project Number SSTF-BA1602-11.

## Author contributions

Conceptualization: H.K., C.R.L., H.J., B.K.K.; Methodology: H.K., B.H.L., C.R.L., H.J.; Software: B.H.L., C.W.L., Y.S.; Formal analysis: C.R.L., H.J., B.H.L., C.W.L., Y.S., J.K.; Investigation: C.R.L., H.J., Y.S.; Resources: J.-i.K., D.I.C.; Writing: C.R.L., B.H.L., H.Y.P., B.K.K.; Supervision: H.Y.P., B.K.K.; Funding acquisition: H.Y.P., B.K.K.

## Method details

### AAV virus production

In all experiments, we used Adeno-Associated Viruses serotype 1/2 that contains capsids of serotypes 1 and 2. The preparation of AAV1/2 was done as described previously^11^. HEK293T cells were cultured in 150 mm culture dishes and grown to ∼60% confluency on the following day. Each dish was cultured in 18 ml of Opti-MEM (Gibco-BRL/Invitrogen, cat# 31985070) after being transfected with plasmids encompassing the vector of interest, AAV2 ITRs, p5E18, p5E18-RXC1, and pAd-F6. Five days after transfection, the supernatant media containing virus particles was transferred to a 50-ml tube and centrifuged at 3,000 rpm for 10 min. The supernatant was slowly added onto a poly-prep chromatography column (Bio-Rad Laboratories, Inc. cat# 731-1550) containing 1 ml of heparin-agarose suspension (Sigma, cat# H6508). To wash the column, 4 ml of Buffer 4-150 (150 mM NaCl, 10 mM citrate, pH 4.0) and 12 ml of Buffer 4-400 (400 mM NaCl, 10 mM citrate, pH 4.0) were used. The AAV1/2 particles were eluted by 4 ml of Buffer 4-1200 (1.2 M NaCl, 10 mM citrate, pH 4.0). The elute was loaded onto the Amicon Ultra-15 filter unit (Millipore, cat# UFC910024) and centrifuged at 4,000 rpm for 20 min. The AAV1/2 particles stuck in filter were eluted by adding 4 ml DPBS and centrifuging 4,000 rpm for 50 min. The titration of virus particles was done by quantitative RT-PCR.

### Stereotaxic surgery

Adult 8-10-week-old WT mice were used for stereotaxic surgery and further experiments. For stereotaxic surgery, mice were anesthetized by i.p. injection of a ketamine/xylazine solution. Virus mixture was injected with a Hamilton syringe, using a 31-gauge needle. The needle was slowly lowered 0.05mm below the injection site for 2 min. After the needle was returned to the injection site, 0.5 μl of the virus mixture was injected at a rate of 0.125 μl/min. Following a 6-min wait for viral dispersion at the injection site and to prevent backflow, the needle was slowly withdrawn from the skull. Stereotaxic coordinates from bregma were: Right hippocampal CA1 (AP: -1.8 / ML: -1.45 / DV: -1.65 from dura), Left hippocampal CA3 (AP: -1.7 / ML: +2.35 / DV: -2.4). The concentrations of injected viruses are described in Table S1.

### Hippocampal window surgery

For *in vivo* imaging experiments, we used 8–11-week-old male WT mice and implanted hippocampal window above dorsal CA1 of mice a week after virus injection surgery. Mice were fixed to a stereotactic frame after being given an intraperitoneal injection of ketamine / xylazine for anesthesia. Chronic hippocampal windows were implanted as described previously^32^. Briefly, a 2.7-mm diameter trephine drill (FST) was used to make a craniotomy at AP -2.0 mm, ML +1.8 mm from the bregma, covering the dorsal hippocampus. The cortical tissue above the CA1 area was carefully removed by aspiration after the dura was properly removed with forceps. By using Meta-Bond (Parkell), a customized stainless-steel cylindrical cannula with a 2.5-mm Ø round glass coverslip (Marienfeld, custom order) attached at the bottom was inserted and cemented to the skull. Chronic cranial windows were implanted over the dorsal CA1 at AP -1.8 mm, ML -1.45 mm from the bregma. The craniotomy was covered with a 2.5-mm spherical glass coverslip and sealed with Meta-Bond. Before the Meta-Bond was cured, a customized stainless-steel head ring was placed around either the cannula or the window and fixed with additional Meta-Bond.

### Contextual fear conditioning and extinction

Each mouse was single caged immediately after the hippocampal window implant surgery. Before day-1 imaging, each mouse was habituated to the anesthesia chamber without isoflurane for 3 minutes on 5 consecutive days. Mice were conditioned on the next day of day-1 imaging. Two hours before conditioning, 100 μg/g doxycycline was injected by i.p. injection during brief anesthesia by isoflurane in the anesthesia chamber. Conditioning sessions were 300 sec in duration, and three 0.75 mA shocks of 2 s duration were delivered at 208 s, 238 s, and 268 s in a square chamber with a steel grid (Med Associates Inc., St Albans, VT). When the contextual fear conditioning was finished, mice were immediately transferred to their homecage. Two days after conditioning, day-2 imaging was performed. On the following day, extinction groups were exposed to the conditioned context for 3 consecutive days, whereas no extinction group stayed at homecage. After day-6 imaging, mice were exposed to the same context to measure freezing levels and were perfused for immunohistochemistry.

### Sample preparation and confocal imaging

Mice were deeply anaesthetized with a ketamine/xylazine solution and perfused with PBS and PBS with 4% paraformaldehyde (PFA). The implanted hippocampal window in the brain was carefully removed, and the brain was further fixed with 4% PFA solution overnight at 4°C. The brain was dehydrated in PBS with 30% sucrose for 2 days at 4°C. The brain was frozen and sliced with a cryostat in 40 μm sections for immunohistochemistry or 50 μm for confocal imaging. Brain slices were mounted with VECTASHIELD mounting medium (Vector Laboratories). For dual-eGRASP imaging, dendrites in CA1 stratum radiatum were imaged in Z-stack with Leica SP8 confocal microscope with 63x distilled water immersion objective lens.

### *In vivo* two-photon imaging

The dorsal CA1 was imaged through the hippocampal window using a two-photon excitation laser scanning microscope (Olympus, FVMPE-RS). The two-photon microscope was equipped with four photomultiplier tubes, a Ti:Sapphire laser (Mai-Tai DeepSee, Spectra-Physics), a galvo/resonant scanner, and a 25× 0.95 NA water immersion objective with an 8-mm working distance (Olympus, XLSLPLN25XSVMP2). Excitation wavelengths of 880 nm (for Cyan GRASP and iRFP imaging) and 960 nm (for Yellow GRASP and mScarlet-i imaging) were used. Cyan GRASP and Yellow GRASP fluorescence were reflected by a long-pass dichroic mirror (Olympus, FV30-SDM570), followed by passing a filter cube (Olympus, FV30-FCY) that consisted of a 505 nm long-pass dichroic mirror, 460-500 nm band-pass filter, and 520-560 nm band-pass filter. mScarlet-i and iRFP signal were collected after passing through a filter cube (Olympus, FV30-FRCY5) that consist of 650 nm long-pass dichroic mirror, 575-645 nm band-pass filter, and 660-750 nm band-pass filter. To reduce motion artifacts, we collected 30 images for each z-plane using a resonant scanner that has a 30 Hz image acquisition rate. In each area, we scanned 84.8 μm× 84.8 μm× 30 μm before fear conditioning and 84.8 μm× 84.8 μm× 32 μm after fear conditioning. During two-photon imaging, mice were anesthetized by isoflurane inhalation using a low-flow vaporizer (Kent Scientific, SomnoSuite), and the body temperature was maintained at 37 °C.

### Image processing

For synapse-level two-photon imaging, it was critical to correct for motion artifacts due to cardiac and respiratory rhythms. 30 images for each z-stack acquired with resonant/galvo scanner were motion corrected with NoRMCorre^33^ package on MATLAB 2021a. After averaging the images at each z-plane, four channel images from 880 nm and 960 nm excitation were aligned by MultiStackReg plugin^34^ on Fiji. Deconvolution of two-photon microscopy images was performed by using the Parallel Iterative Deconvolution plugin on Fiji.

### IMARIS analysis

Imaris (Bitplane, Zurich, Switzerland) software was used to process and analyze the two-photon images. Each trackable myr_mScarlet-I-positive or myr_iRFP670-only dendrites were manually denoted as engram or non-engram dendrites by a researcher, while other researchers were blinded from the information during analysis. Each cyan or yellow eGRASP signal was manually denoted with the annotation tool within Imaris. When the cyan and yellow eGRASP signals overlapped in a single synapse, it was denoted as overlapping annotations as the presynaptic neuron of the synapse indicating IEG-positive during memory formation. The length of each dendrite on day-1 and day+2 was calculated using Imaris measurement and was averaged. For relative density analysis, cyan and yellow eGRASP density of each dendrite was normalized to the average spine density of the cyan and yellow eGRASP on the myr_iRFP670-only dendrites, respectively. In all IMARIS analysis, the investigators who analyzed the images were blinded to the behavior group of mice.

### Spine distribution analysis

For spine distribution analysis, coordinates of all denoted spots and their labels were used to calculate distance between spines in the custom MATLAB code. To evaluate the density of surrounding spines in Fig. 4D, we first calculated the first principal component vector using the coordinate of all spines on the dendrite segment. By projecting the spines on the principal vector line, we generated a one-dimensional spine map. To calculate the density of spine, we binned the spine location every 0.06 μm, and filtered it with a gaussian window of 2.5 μm standard deviation.

### Statistics

Data analysis and figures were prepared using Prism software. To analyze GRASP data that was not normally distributed, the Mann-Whitney test was used. The value of n and statistical significance were described in each figure legends.

### Materials & Correspondence

Correspondence and material requests should be addressed to Bong-Kiun Kaang and Hye Yoon Park.

## Supplemental Figures

**Figure S1. Total cyan GRASP synaptic density across different memory states**.

Synaptic densities of cyan GRASP on, before and after the fear conditioning or extinction. Each data point in the “Before” graph represents a dendrite. The same dendrite is represented in the “After” graph. The densities of “cyan only” or yellow puncta on engram dendrites are raw values (the number of puncta/μm). (A) Absolute density of T-N synapses. CA1 non-engram dendrites (n = 35); Mann-Whitney two-tailed test. Dunn’s multiple comparisons test after Kruskal-Wallis test. Kruskal-Wallis: ****P < 0.0001; Dunn’s ***P < 0.001, ****P < 0.0001. (B) Absolute density of T-E synapses. CA1 engram dendrites (n = 34); Mann-Whitney two-tailed test. Dunn’s multiple comparisons test after Kruskal-Wallis test. Kruskal-Wallis: ****P < 0.0001; Dunn’s ***P < 0.001, ****P < 0.0001. Data are represented as mean ± SEM in all figures.

**Figure S2. Freezing levels of mice across contextual fear conditioning and extinction**.

(A) Experimental scheme of CFC and fear extinction paradigm. Extinction group (N = 4) was exposed to the conditioning context on three consecutive days of extinction session while no extinction group (N = 3) stayed in their home cages. (B) Freezing levels of mice measured before and after contextual fear conditioning. N = 5, Unpaired t-test, *P = 0.0156, t_8_ = 3.058 (C) Freezing levels of extinction (N = 4) or no extinction group (N = 3). Mice that did not show effective fear extinction (freezing level higher than 50% during fear retrieval) were excluded from analysis. Unpaired t-test, *P = 0.0107, t_5_ = 3.965 (D) Trace of freezing levels across fear extinction sessions. Basal freezing levels were measured for 5 minutes during a fear conditioning session. Freezing levels during extinction were measured every 5 minutes in each session. (Extinction group, N = 4). Data are represented as mean ± SEM in all figures.

**Figure S3. Relative density and proportions of synapses in context only group**.

(A) Schematic illustration of the experimental protocol to examine synaptic connections through context exposure. (B) Relative density of random cyan GRASP on non-activated or activated dendrites before and after context exposure; CA1 non-activated dendrites (n = 43); CA1 activated dendrites (n = 14); Mann-Whitney two-tailed test. (Left, Before CFC) **P = 0.0090. (Right, After CFC) P = 0.2208. N_0_ and A_0_ indicate CA1 cells before CFC that become non-activated and activated cells later after CFC, respectively. (C) Relative density of yellow GRASP on non-activated or activated dendrites after context exposure; CA1 non-activated dendrites (n = 43); CA1 activated dendrites (n = 14); Mann-Whitney two-tailed test. P = 0.0947. (D) The absolute densities of “existing” or “newly formed” synapses. CA1 non-activated dendrites (n = 43); CA1 activated dendrites (n = 14); (Left) Mann-Whitney two-tailed test. P = 0.0884; (Right) Mann-Whitney two-tailed test. P > 0.9999. (E) The proportion of “existing” and “newly formed” synapses in non-activated or activated dendrites; “existing” N-A synapses (n = 114); “new” N-A synapses (n = 1); “existing” E-E synapses (n = 75); “new” E-E synapses (n = 2); CA1 non-activated dendrites (n = 43); CA1 activated dendrites (n = 14). (F) Proportions of persisting and disappearing synapses between activated cells in the context only group. Yellow and gray indicate persisting and disappeared percentages of the synapses, respectively. Disappeared “existing” A-A synapses (n = 2); persisting “existing” A-A synapses (n = 70); persisting “new” A-A synapses (n = 2); disappeared “existing” A-N synapses (n = 2); persisting “existing” A-N synapses (n = 113); persisting “new” A-N synapses (n = 1). (G) Absolute synaptic density of existing and new A-A synapses in the context only group. CA1 engram dendrites (n = 43); CA1 non-engram dendrites (n = 14). (H) Absolute synaptic density of existing and new A-N synapses in the context only group. CA1 engram dendrites (n = 43); CA1 non-engram dendrites (n = 14).

**Figure S4. Normalized fraction of disappeared and persisting synapses of extinction and no extinction group**.

Fraction of disappeared synapses after fear extinction is significantly higher than no extinction group only in E-E synapses, while E-N synapses show insignificant difference. Two-sided Chi-square test, χ^2^(1) = 18.85, ****P < 0.0001 (‘Existing’ E-E synapses of extinction and no extinction groups); χ^2^(1) = 4.177, *P = 0.0410 (New E-E synapses of extinction and no extinction groups); χ^2^(1) = 1.278, P = 0.2583 (Existing E-N synapses of extinction and no extinction groups); χ^2^(1) = 0.4958, P = 0.4813 (New E-N synapses of extinction and no extinction groups).

