## Supplementary figures and images for "Hippocampal engram networks for fear memory recruit new synapses and modify pre-existing synapses *in vivo*"

### Supplementary figure 1

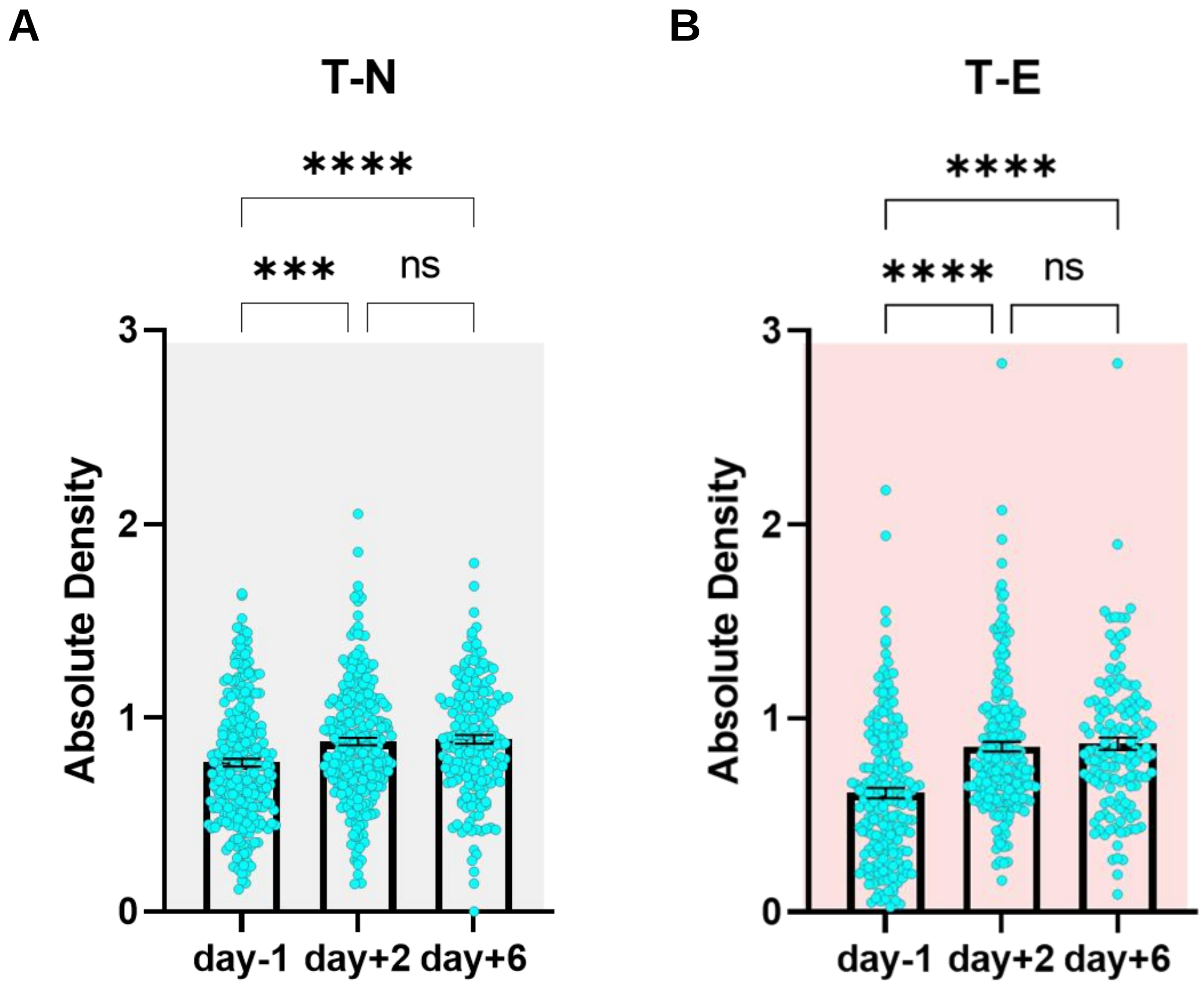

### Supplementary figure 2

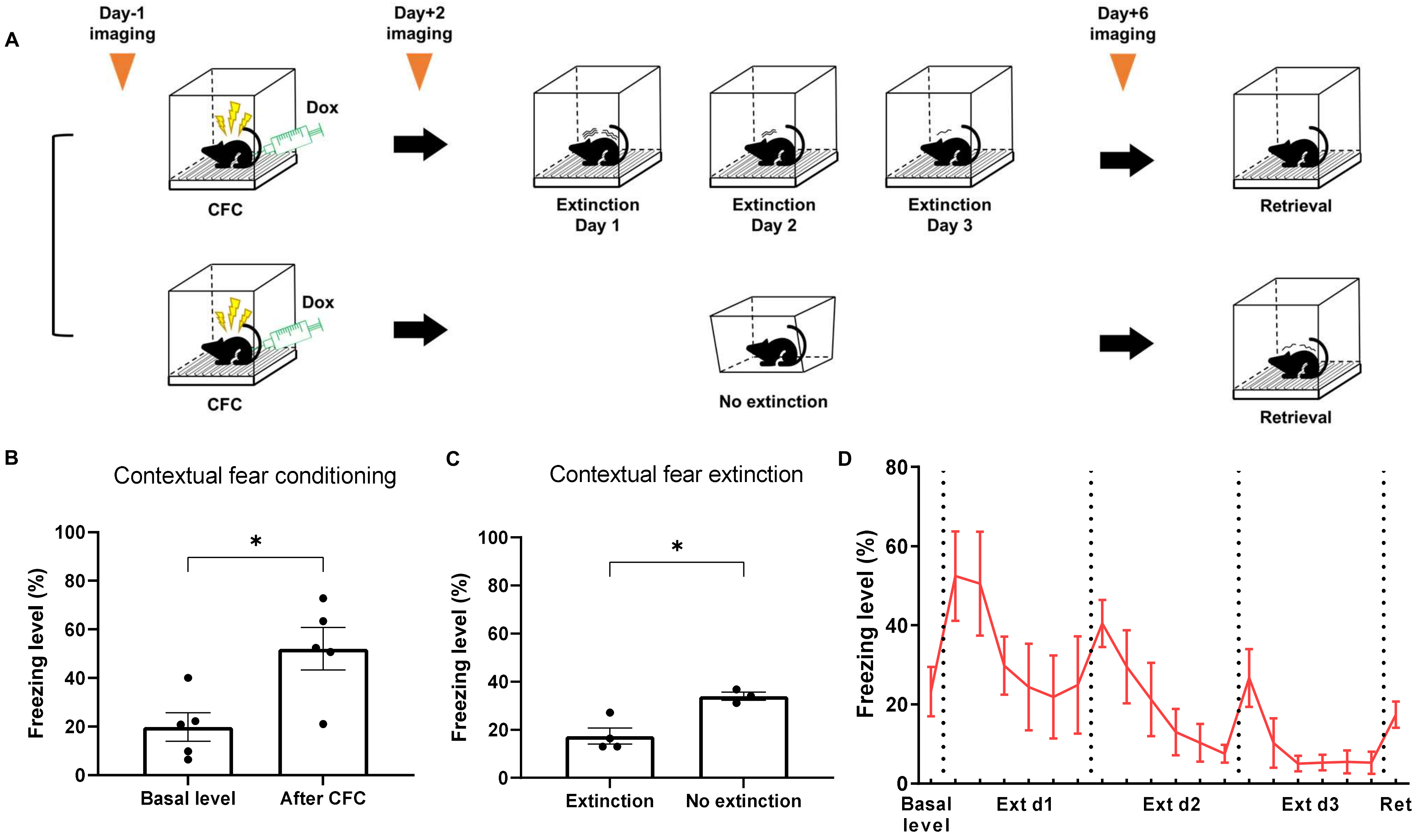

### Supplementary figure 3

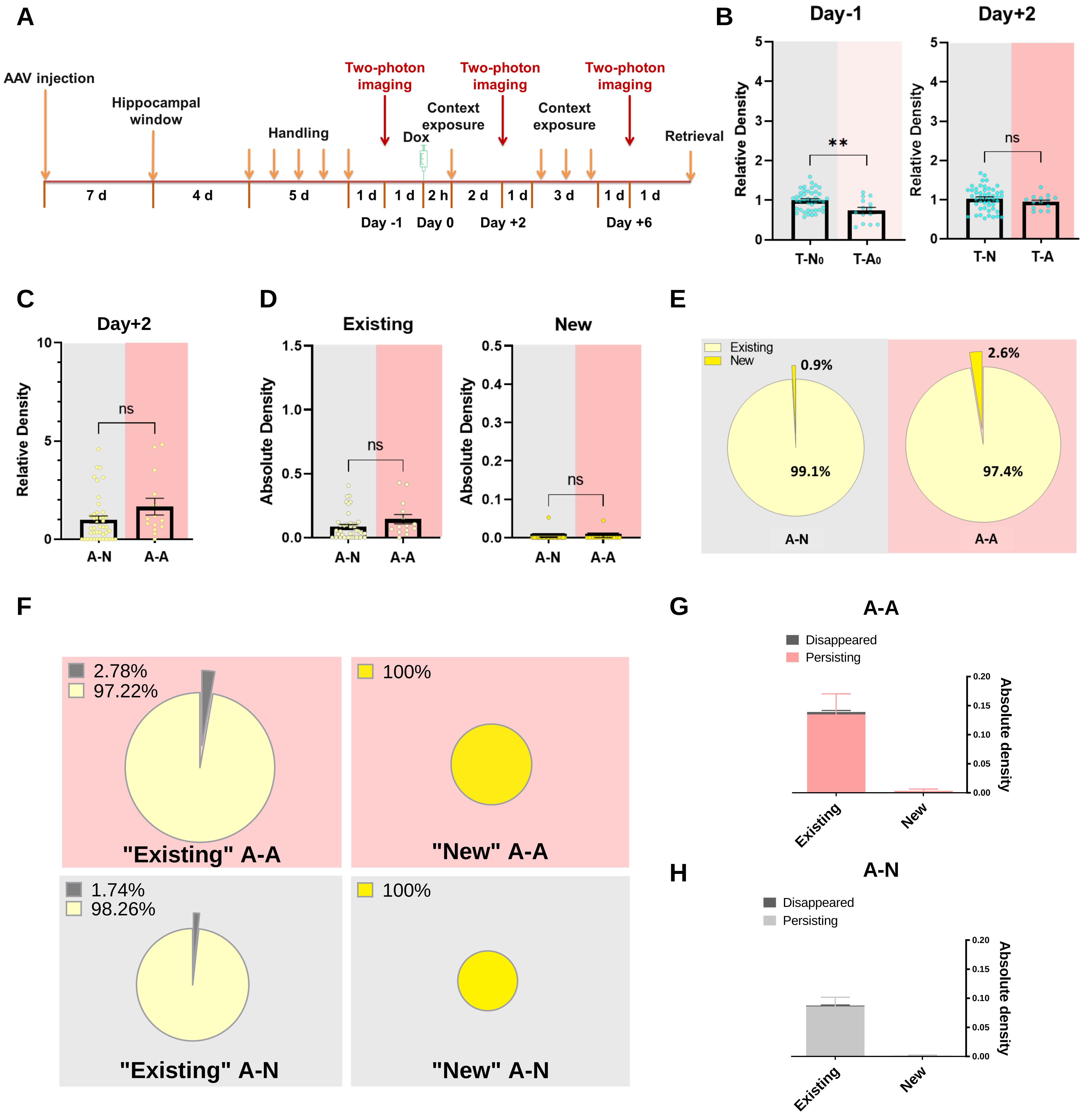

### Supplementary figure 4

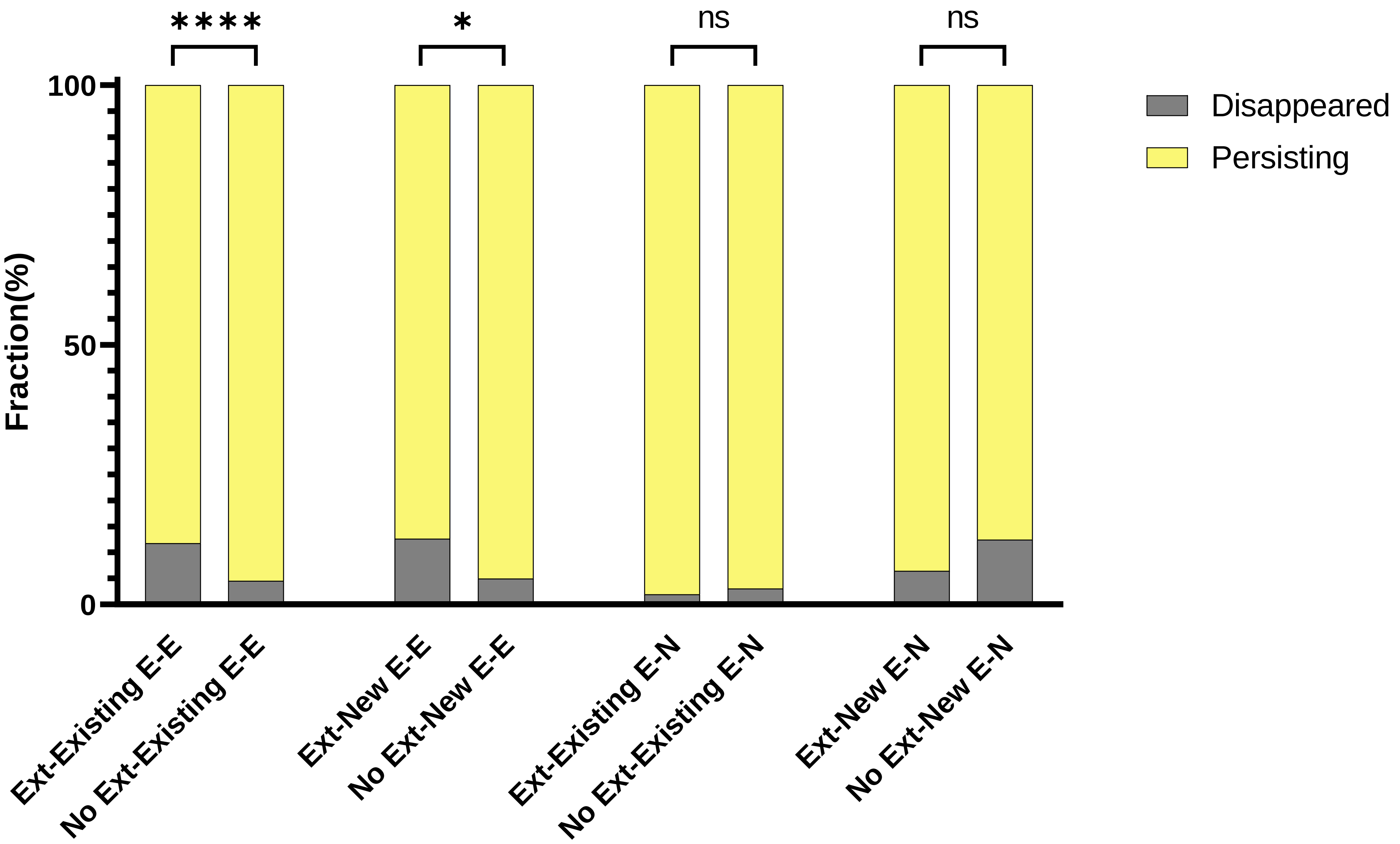
