## Supplementary table 1 for "Hippocampal engram networks for fear memory recruit new synapses and modify pre-existing synapses *in vivo*"

**Table S1.** Virus volumes and concentrations used for experiments, related to STAR Methods.

| **Region** | **Virus** | **Volume per injection** | **Stock titer** | **Source** |
| --- | --- | --- | --- | --- |
| CA1 | AAV2/1-Fos-rtTA | 0.23 µL | 2.59e9 vg/µL | In-house |
|  | AAV2/1-CamKll-iCre | 0.09 µL | 2.63e6 vg/µL | In-house |
|  | AAV2/1-EF1a-DIO-myriRFP670V5-P2A-post-eGRASP | 0.41 µL | 8.85e9 vg/µL | In-house |
|  | AAV2/1-TRE3G-myrmScarlet-I | 0.27 µL | 7.41e9 vg/µL | In-house |
| CA3 | AAV2/1-Fos-rtTA | 0.19 µL | 2.59e9 vg/µL | In-house |
|  | AAV2/1-CamKll-cyan pre-eGRASP | 0.45 µL | 3.11e9 vg/µL | In-house |
|  | AAV2/1-TRE3G-yellow pre-eGRASP | 0.35 µL | 4.50e9 vg/µL | In-house |
