## Supplementary material for "Hippocampal engram networks for fear memory recruit new synapses and modify pre-existing synapses *in vivo*": Key resource table

**Key resources table**

| REAGENT or RESOURCE | SOURCE | IDENTIFIER |
| --- | --- | --- |
| Bacterial and virus strains | | |
| AAV2/1-Fos-rtTA | Choi et al., 2018 | N/A |
| AAV2/1-CamKll-iCre | Choi et al., 2018 | N/A |
| AAV2/1-EF1a-DIO-myriRFP670V5-P2A-post-eGRASP | Choi et al., 2018 | N/A |
| AAV2/1-TRE3G-myrmScarlet-I | This paper | N/A |
| AAV2/1-CamKll-cyan pre-eGRASP | Choi et al., 2018 | N/A |
| AAV2/1-TRE3G-yellow pre-eGRASP | Choi et al., 2018 | N/A |
| Chemicals, peptides, and recombinant proteins | | |
| Heparin-Agarose | Sigma-Aldrich | H6508 |
| Doxycycline hyclate | Sigma-Aldrich | D9891 |
| Experimental models: Organisms/strains | | |
| Mouse: C57BL/6 N | Samtako. Bio. Korea | N/A |
| Recombinant DNA | | |
| pAAV-Fos-rtTA | Choi et al., 2018 | Addgene 120309 |
| pAAV-CamKll-iCre | Choi et al., 2018 | N/A |
| pAAV-EF1a-DIO-myriRFP670V5-P2A-post-eGRASP | Choi et al., 2018 | Addgene 111585 |
| pAAV-TRE3G-myrmScarlet-I | This paper | N/A |
| pAAV-CamKll-cyan pre-eGRASP | Choi et al., 2018 | Addgene 111586 |
| pAAV-TRE3G-yellow pre-eGRASP | Choi et al., 2018 | N/A |
| Software and algorithms | | |
| IMARIS | Bitplane | https://imaris.oxinst.com/ |
| Prism 9 | GraphPad | https://www.graphpad.com/ |
| MATLAB | MathWorks | https://kr.mathworks.com/ |
| Video Freeze® | Med Associates Inc® | https://www.med-associates.com/ |
| FV30S-SW | Olympus | https://www.olympus-lifescience.com/ |
| FV30S-DT | Olympus | https://www.olympus-lifescience.com/ |
| ImageJ | Schneider et al., 2012 | https://imagej.nih.gov/ij/ |
| Other | | |
| FVMPE-RS (two-photon excitation laser scanning microscope) | Olympus | https://www.olympus-lifescience.com/ |
